# Clusterization in head and neck squamous carcinomas based on lncRNA expression: molecular and clinical correlates

**DOI:** 10.1101/105999

**Authors:** Pelayo G. de Lena, Abel Paz-Gallardo, Jesús M. Paramio, Ramón García-Escudero

## Abstract

**Background:** Long non-coding RNAs (lncRNAs) have emerged as key players in a remarkably variety of biological processes and pathologic conditions, including cancer. Next-generation sequencing technologies and bioinformatics procedures predict the existence of tens of thousands of lncRNAs, from which we know the functions of only a handful of them, and very little is known in cancer types such as head and neck squamous cell carcinomas (HNSCCs).

**Results:** Here, we use RNA-seq expression data from The Cancer Genome Atlas (TCGA) and various statistic and software tools in order to get insight about the lncRNome in HNSCC. Based on lncRNAs expression across 426 samples, we discover five distinct tumor clusters that we compare with reported clusters based on various genomic/genetic features. Results demonstrate significant associations between lncRNA-based clustering and DNA-methylation, TP53 mutation, and human papillomavirus infection. Using “guilt by association” procedures, we infer the possible biological functions of representative lncRNAs of each cluster. Furthermore, we found that lncRNA clustering is correlated with some important clinical and pathologic features, including patient survival after treatment, tumor grade or sub-anatomical location.

**Conclusions:** We present a landscape of lncRNAs in HNSCC, and provide associations with important genotypic and phenotypic features that may help to understand the disease.

## Background

Head and neck cancer is the sixth leading cancer worldwide, with an estimated 600,000 new cases annually and a 50% five-year mortality rate (Globocan 2012) [1]. As more than 90% of this cancer cases are of squamous origin, they are generally referred to as head and neck squamous cell carcinoma (HNSCC). HNSCC arises in the upper aerodigestive tract, comprising the nasal cavity and paranasal sinuses, oral cavity, pharynx, larynx, and trachea. The main risk factors associated with its development are tobacco and alcohol consumption, which have a synergistic effect when combined, and also human papillomavirus (HPV) infection [2, 3]. HPV is known to drive tumorigenesis through the actions of its major oncoproteins E6 and E7, which can inactivate p53 and retinoblastoma (Rb) tumor supressors, respectively, altering cell cycle regulation in infected cells. HPV-positive differ from HPV-negative HNSCCs in tumor biology and clinical characteristics, including clinical outcomes, since HPV-positive tumors have been associated with a more favorable prognosis [4]. HNSCC patients are frequently treated with surgery, together with radiotherapy and/or cisplatin-based chemotherapy. Patients with aggressive disease are treated with cetuximab, an anti-EGFR antibody. Few patients respond to this therapy, and there is not molecular stratification of the patients neither biomarkers of responsiveness [2]. Previous characterization of molecular features in HNSCC, particularly with the aid of large-scale cancer genomics initiatives such as the TCGA, has generated important insights for stratifying patients and delineating tumor subtypes [5-7]. These multiomic analyses, do not take into account the vast long non-coding transcriptome that may substantially contribute to HNSCC pathogenesis and progression.

LncRNAs are transcripts longer than 200 nucleotides that have no apparent protein coding potential [8]. They are highly diverse and actively present in many aspects of cell biology, including cellular differentiation, proliferation, DNA damage response, dosage compensation, and chromosomal imprinting. LncRNAs are categorized as exonic, intronic, intergenic, antisense, or overlapping based on their genomic location relative to a protein-coding gene. The most recent estimate of the Encyclopedia of DNA Elements (ENCODE) Project Consortium (GENCODE version 25) is that the human genome contains more than 15,000 lncRNA genes that encode almost 28,000 transcripts [9] although the total number is estimated to be much higher. Like the protein coding genes (PCGs), lncRNAs genes are regulated transcriptionally and by histone modification, and lncRNA transcripts are processed by the canonical splicing machinery [10]. In addition, lncRNAs have fewer exons than PCGs, are usually located in the nucleus, are subject to less selective pressure during evolution, and show higher tissue or cell type specificity. Given the large number of lncRNAs that are predicted to exist, it is expected that the functions in which they are involved may include many of the known (and possibly new) biological and physiological processes. Some of the molecular functions described so far include chromatin interactions, transcriptional regulation, RNA processing, mRNA stability or translation, or signaling cascades modulation [11]. In the context of cancer phenotypes, lncRNAs functions have been found in proliferation, growth suppression, motility, immortality, angiogenesis and viability [11, 12]. Unfortunately, the possible functions or phenotypic effect of the vast majority of lncRNAs remains elusive in normal homeostasis or in cancer [12, 13], and are difficult to analyze [14].

Comprehensive genomic analysis across human cancers demonstrated that a large number of lncRNAs show differential expression among known tumor subtypes [15-17]. In addition, lncRNA alterations are highly tumor and lineage specific [15-17]. The expression of known, tumor associated lncRNAs, such as HOTAIR, NEAT1, UCA1, MALAT1, and MEG3, has been tested in HNSCC and correlated with clinicopathologic parameters [18]. Functional studies have tested the effects in proliferation, apoptosis, invasion and migration in HNSCC cell lines after RNA interference (RNAi) of specific lncRNAs [19]. In addition, lncRNA profiling has been done in HNSCC, assessing deregulation between normal and tumor samples, associations with clinical parameters, HPV infection or mutation in the TP53 tumor suppressor gene [19-21].

To our knowledge, no comprehensive reports studying the clusterization of human primary HNSCC samples based on genome-wide lncRNAs expression have been published. Therefore, the main objective of this report is to analyze the long non-coding transcriptome in HNSCC in order to discover new tumor clusters. Furthermore, we investigate whether lncRNA-based clusterization is useful to predict patient’s clinical outcome, and is associated with important clinicopathologic parameters. Finally, we interrogate the dataset to infer the possible biological functions of lncRNAs in HNSCC based on correlation patterns of the PCGs.

## Methods

### Data resources

Expression values of 12,727 lncRNA genes from 426 HNSCC primary tumor samples of the TCGA RNAseq cohort were downloaded from the TANRIC web page [22]. Methods used to extract these expression values have been described [16]. Briefly, the genomic coordinates of the human lncRNAs from the GENCODE Resource (version 19) were obtained. Thereafter, the lncRNA exons that overlapped with any known coding genes based on the gene annotations of GENCODE and RefGene were filtered out. As a result, the analysis focused on the remaining 12,727 lncRNAs. Based on the BAM files, the expression levels were quantified as RPKM, and the lncRNAs with detectable expression were defined as those with an average RPKM ≥ 0.3 across all samples in each cancer type, as defined in the literature. The Ensembl identifiers of the 500 lncRNA genes displaying the highest variability in HNSCC were kindly provided by Dr. H. Liang at the Research Group from the Department of Bioinformatics and Computational Biology of The University of Texas and the MD Anderson Cancer Center. We downloaded DNA methylation, CNV, miRNA, and reverse phase protein array (RPPA) clustering data, as well as HPV infection and PCG mutations for the TCGA HNSCC dataset using the cBioPortal [23], the UCSC Xena [24] and the TCGA [25] repositories.

### Unsupervised clustering analysis

Consensus Cluster Plus (CCP) tool [26], which is implemented as an R language package from Bioconductor [27], extends the CC algorithm and is briefly described here. The algorithm begins by subsampling a proportion of items and a proportion of features from a data matrix. Each subsample is then partitioned into up to k groups by a user-specified clustering algorithm. This process is repeated for a specified number of times. Pairwise consensus values, defined as ‘the proportion of clustering runs in which two items are grouped together’, are calculated and stored in a consensus matrix (CM) for each k. Clustering settings used were as follows: maxK=6; number of bootstraps=1000; item subsampling proportion=0.8; feature subsampling proportion=1; cluster algorithm=pam; inner linkage type=complete; final linkage type=complete; correlation method=Euclidean. Consensus CDF plot and proportion of ambiguous clustering (PAC) per each k was obtained.

### Analysis of tumor clusters revealed by lncRNA expression

The association between lncRNA-based sample clusters and sample clusters based on other molecular features/aberrations was done using Fisher’s exact test for fourfold (2x2) tables or Chi-square test for more than fourfold comparisons. Both tests are used to determine whether there are significant differences between the expected frequencies and the observed frequencies in one or more categories. Kaplan-Meier curves were obtained using the follow-up time of the HNSCC patients of 2 end-points: recurrence and death. Statistical analyses were done with SPSS 14.0.

### Guilt-by-association (GBA) analysis

Pearson correlation analysis was used to select lncRNAs genes with significant correlation between artificial expression vectors or templates (t1 to t5) (Fig. 2A), or between mean expression values of correlated lncRNAs and PCGs. We select both direct and inverse correlation patterns by setting thresholds either at the Pearson (r) value or at the associated p-val. Pearson’s r values ranges between −1 to +1, such that 2 perfectly correlated genes display r=1, and 2 perfectly anticorrelated genes r=-1. LncRNAs surrogate selection was done setting r>0.3 with respect to the corresponding template (associated p-val<1×10^−6^). Additional filtering criteria include average RPKM ≥ 0.1 within specific clusters, and overexpression between normal tissue and specific clusters (Ttest, corrected p-val<0.05). PCGs directly or inversely correlated with lncRNAs were selected using r>0.3, or r<-0.3 with respect to mean expression values of selected lncRNAs per cluster, respectively (associated p-val<1×10^−9^). Correlations as well as heatmap drawings were performed using MultiExperiment Viewer v4.9 (MeV) [28].

### Gene Ontology analysis

Selected PCGs were analyzed with the Gene Functional Annotation Tool available at the DAVID v6.7 website [29, 30] using their official gene symbols. Gene ontology option GOTERM_BP_FAT was selected and a functional annotation chart generated. A maximum p-value of 0.05 was chosen to select only significant categories.

## Results

### HNSCC sample clusters based on lncRNA expression

In order to discover HNSCC tumor subgroups, we selected the 500 lncRNAs (Supplementary Table 1) with the most variable expression pattern in HNSCC [16] and the Consensus Cluster Plus (CCP) software tool (see Methods). CCP analysis revealed the presence of five HNSCC clusters, which we have named clusters 1 to 5 (Fig. 1 and Supplementary Table 2). The results suggest that lncRNAs can distinguish 5 HNSCC subtypes, having significant differences in lncRNA expression.

**Figure 1.**
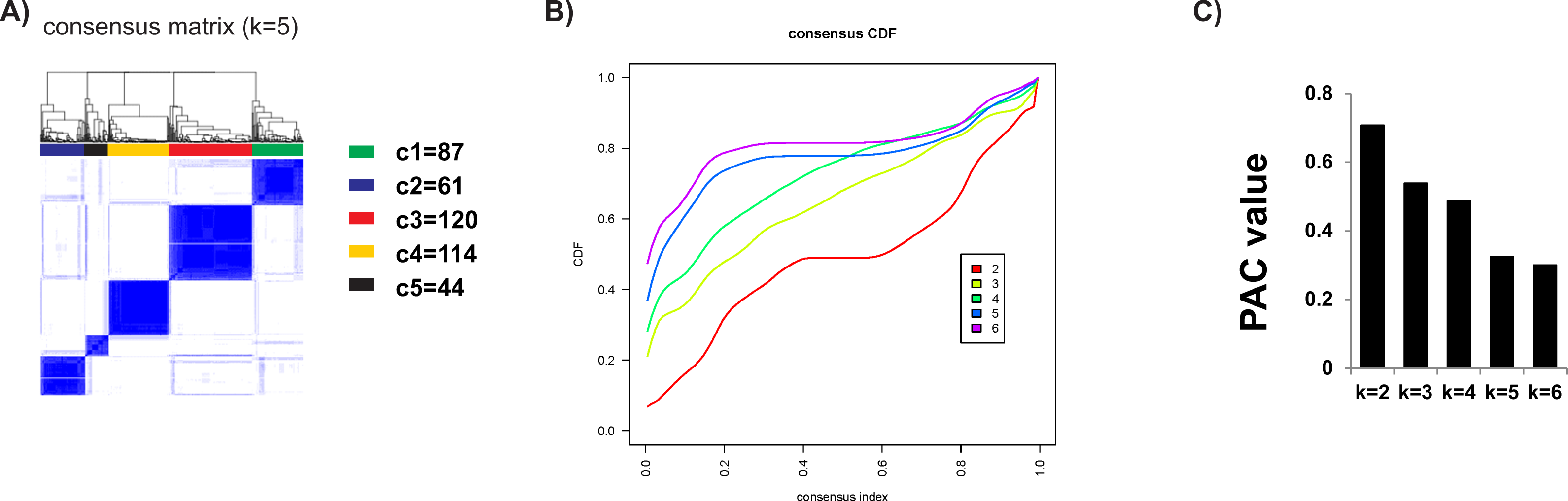
Unsupervised clustering of HNSCC using lncRNA expression data. **A)** Consensus Cluster Plus analysis identifies five major groups (samples, n=426). The blue and white heat map displays sample consensus. Number of samples per cluster is shown. Consensus CDF plot **(B)** and PAC values **(C)** for k=2 to 6 are represented. Smaller PAC values are obtained with k=5 and k=6, with minor differences between them. Therefore, k=5 was selected. Specifications and parameters used in the analysis are described in the Methods section.

### LncRNAs clustering resembles DNA methylation and mRNA clustering

In order to assess the correlation between lncRNAs and additional molecular features, we analyze whether lncRNA clustering resembles previously described sample clustering based on mRNA, miRNA, protein (RPPA) expression, DNA methylation and CNV. For this, we downloaded clustering data from the TCGA-HNSSC dataset [5] and compared them with lncRNAs subgroups using contingency analyses (Chi-square test, see Methods). The results showed highly significant similarities between our clusterization and DNA methylation clusters (p-val=4.6 × 10^−34^) or mRNA-based subtypes (p-val=4.2 × 10^−30^) (Fig. 2A). LncRNA clustering displays lower overlapping with CNV, miRNA, and RPPA subtypes, with p-values ranging from 2.8 × 10^−15^ to 3.5 × 10^−5^ (Fig. 2A). The high overlapping between lncRNA and mRNA subgroups is suggestive of similar molecular mechanisms of expression control. Also, the contingency results suggest that lncRNA expression might be strongly influenced by DNA methylation (or viceversa), an important mechanism of epigenetic transcriptional regulation.

**Figure 2:**
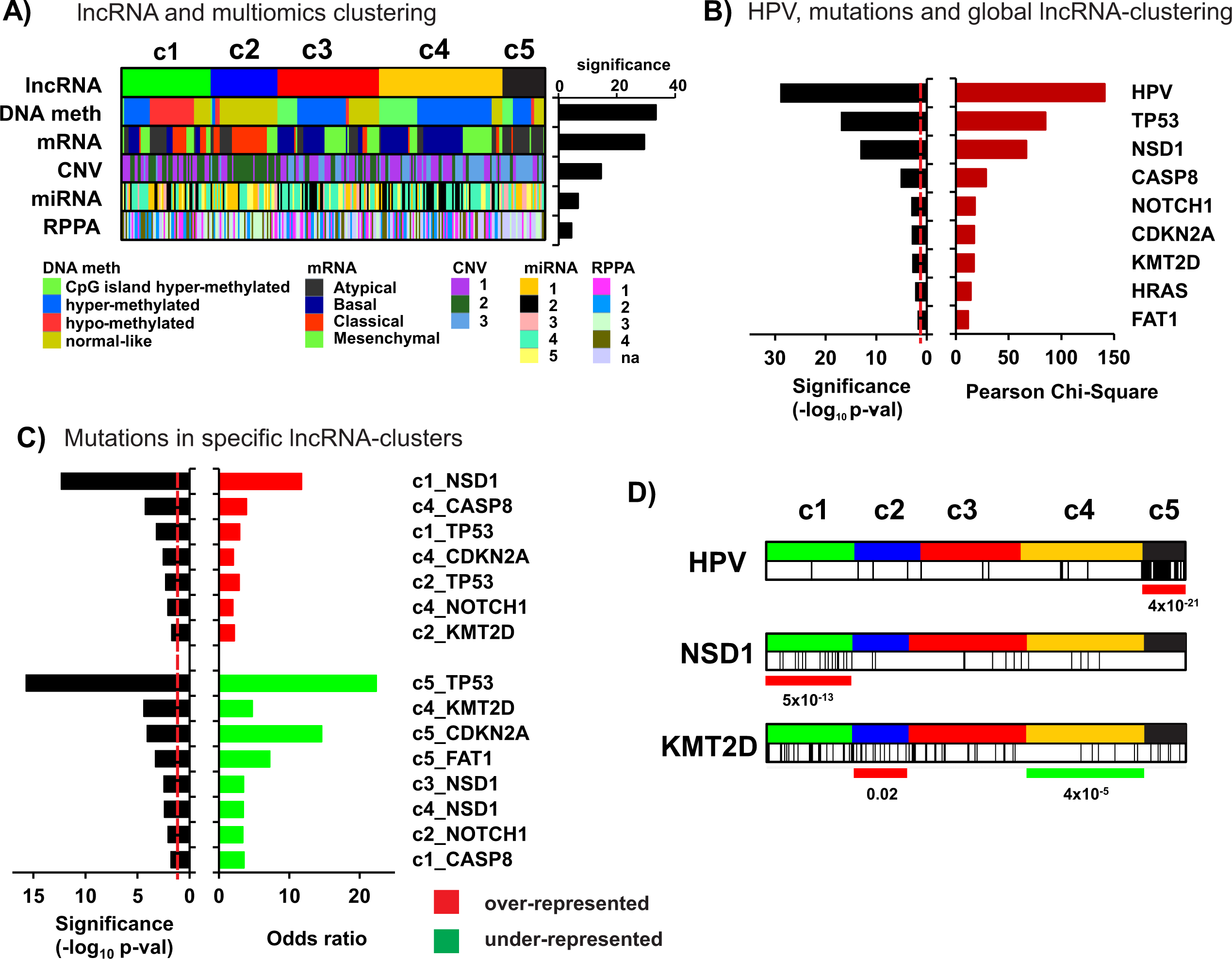
lncRNA clusters and other molecular aberrations. **A)** LncRNA-based clustering of HNSCC samples is significantly associated with clustering based in diverse molecular features, mainly DNA methylation and expression of PCGs (mRNA). Significance values are plot upon Chi-square test computation. **B)** and **C)** Association with HPV infection and HNSCC mutations with lncRNA clusters. Chi-square or odds ratio values are plot upon Chi-square test **(B)** or Fisher’s exact test **(C)** computation, respectively. Dashed red line: threshold of significance (p-val<0.05). **D)** Distribution of lncRNA clusters and HPV infected samples, or samples with mutations in KMT2D or NSD1. Note the enrichment of HPV infection in c5, the NSD1 mutations in c1 and the KMT2D mutations in c2 (red lines), and the depletion of KMT2D mutations in c4 (green line). P-values are calculated with Fisher’s exact test. Vertical black lines in panel D showed HPV+ samples and mutated samples for the selected genes KMT2D and NSD1, respectively.

### HPV infected tumors display and specific lncRNAome

A significant proportion of tumor samples from the reported TCGA dataset [5] having 279 samples, is infected with HPV (almost 13%). Interestingly, a deep, multiplatform analysis of the molecular features of HNSCC primary tumors from the TCGA, demonstrated strong differences between HPV-positive and HPV-negative samples [5]. Infected cancers display different mutational landscape, mainly characterized by the absence of TP53 gene mutations. Chi-square test demonstrated that lncRNA clustering is highly significantly associated with HPV (p-val=1.5 × 10^−29^) (Fig. 2B), indicating infected tumors display a specific lncRNA landscape. Interestingly, a deeper analysis shows that almost all samples from cluster 5 (c5) are HPV positive (24 out of 29) (p-val=4.0 × 10^−21^) (Fig. 2D). Similar results were reported in the TANRIC study [16].

### HNSCC mutations and lncRNA clustering

In order to discover whether particular lncRNA clusters are characterized by the presence or absence of HNSCC mutations, we interrogate lncRNA-cluster and mutation associations using Chi-square test (Fig 2B-D). We selected a total number of 19 genes, found to be significantly mutated in the TCGA-HNSCC cohort: CDKN2A, FAT1, TP53, CASP8, AJUBA, PIK3CA, NOTCH1, KMT2D, NSD1, HLA-A, TGFBR2, HRAS, FBXW7, RB1, PIK3R1, TRAF3, NFE2L2, CUL3 and PTEN. Significant associations between these mutations and lncRNA-based clustering are shown (Fig. 2C). Cluster 1 (c1) is enriched in TP53 and NSD1 mutations, cluster 2 (c2) display frequent TP53 and KMT2D mutations, and cluster 4 (c4) frequent CASP8, NOTCH1 and CDKN2A mutations. In addition, cluster c5 is depleted of TP53 and CDKN2A mutations, as most samples are HPV-infected. Finally, cluster 4 (c4) is depleted on KMT2D mutations. Both KMT2D and NSD1 are methyltransferases involved in methylation of H3K4 and H3K36, respectively, which are important epigenetic regulators of gene expression. Most mutations found for both proteins are truncating, therefore indicating that c1 and c2 are characterized by loss-of-function of histone modifiers.

### Guilt-by-association (GBA) analysis: selection of surrogate lncRNAs per cluster

In order to select candidate lncRNAs for further analysis, we decided to perform differential expression analysis using the clustering information. Our aim was to select lncRNAs surrogate of each of the 5 clusters, so they are overexpressed only in 1 cluster. We designed 5 expression vectors or templates, (t1 to t5, see Figure 3A) from which we performed Pearson correlation analysis (see Methods) onto the lncRNA genes from which expression values are available for HNSCC (n=12,727). The output are lncRNAs whose expression patterns are i) expressed in the specific cluster, ii) overexpressed in the specific cluster versus normal tissue, and iii) significantly correlated with the corresponding template (r>0.3 and p-val<10^−6^). The number of selected lncRNAs per cluster and the heatmap showing expression patterns is shown (Fig. 3B and Supplementary Table 3): c1=91, c2=275, c3=9, c4=13, and c5=245. Note that clusters c2 and c5 display the highest numbers of lncRNA genes, and c3 and c4 the lowest. Interestingly, 16 out of 245 lncRNAs in cluster c5, depleted in TP53 mutations (Fig. 2C and D), have been shown to be overexpressed in TP53 wild type HNSCC samples [20]. In addition, 12 out of 245 c5 lncRNA genes were found upregulated in HPV infected samples [20], in line with high frequency of HPV positive samples in this cluster (Fig. 2D and Supplementary Table 5).

**Figure 3.**
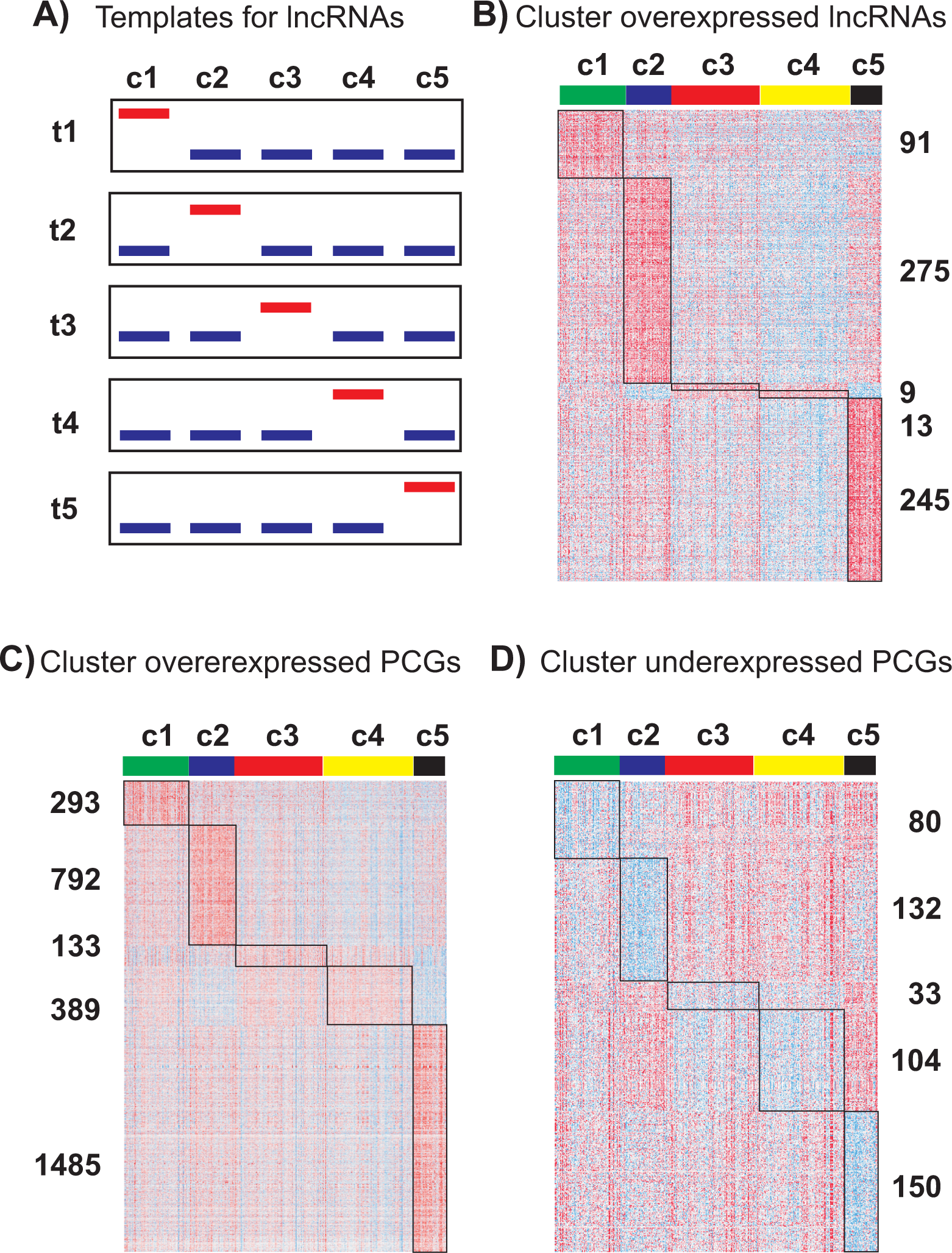
Guilt-by-association (GBA) analysis. **A)** Schema-graph showing approach used to search for lncRNA genes as surrogates of each cluster. Briefly, Pearson correlation was computed between all lncRNAs and artificial expression vectors or templates (templates t1 to t5) whereby maximum expression in individual clusters with respect to the others was interrogated. Threshold used: r>0.3 and p-val<1×10^−6^. Additional filtering criteria include average RPKM ≥ 0.1 within specific clusters, and overexpression between normal tissue and specific clusters (Ttest, corrected p-val<0.05). **B)** Heatmap exhibiting selected lncRNAs upon Pearson correlation and the corresponding gene numbers per cluster. Heatmaps of PCGs directly (overexpressed PCGs) **(C)** or inversely (underexpressed PCGs) **(D)** correlated with the lncRNAs selected per cluster in panel B. Threshold used: r>0.3 and p-val<1×10^−9^ for overexpressed PCGs; r<-0.3 and p-val<1×10^−9^ for underexpressed PCGs.

### Associated PCGs to surrogate lncRNAs

Predicting the biological functions of lncRNAs is challenging. Guilt-by-association (GBA) analysis has been proposed on the basis that the function of a poorly characterized lncRNA gene can be inferred from known/predicted functions of protein coding genes which are co-expressed [31]. We leveraged our RNA-sequencing data using GBA analysis to generate hypotheses on functional significance by comparing the expression of lncRNAs to protein coding genes of known function. Therefore, we performed correlation analysis to interrogate what coding genes are directly or inversely correlated with the surrogate lncRNA genes of each cluster. Accordingly, we use 5 different expression vectors or templates, one per cluster, calculated from the mean expression value of lncRNAs selected per cluster above (Fig. 3B). The number of correlated genes per cluster and the heatmap showing their expression patterns is shown (Fig. 3C and D) for directly (overexpressed PCGs) or inversely (underexpressed PCGs) correlated PCGs. Thresholds used are r>0.3 or r<-0.3 and p-val<1×10^−9^ (Supplementary Table 4). The results shown that each lncRNA cluster have associated a significant number of coding genes, whose functions might be related with the biological roles of the correlated lncRNAs.

### Gene Ontology analysis of lncRNA-clustering associated PCGs

In order to predict possible functions of the surrogate lncRNAs in each cluster, we perform enrichment analysis of GO Biological Processes (GOBP) on the associated PCGs, which would find enriched pathways and larger processes made up of the activities of the PGC gene products. Therefore, we analyze individually the lists of PCGs directly or inversely correlated with each lncRNA cluster (Fig. 4). Cluster c1 is characterized by the underexpression of PCGs involved in T cell, natural killer cell and myeloid leukocyte activation; these processes are overexpressed in c5, suggesting important differences in immune response between c1 and c5. Importantly, PD-L1 (CD274), an immune inhibitory receptor ligand whose inactivation using antibodies is being currently used for cancer treatment successfully, is underexpressed in c1. Cluster c2 is characterized by the expression of positive regulators of transcription, tissue morphogenesis, and neuronal markers. In addition, c2 exhibits depletion of PCGs involved in ubiquitin-dependent protein cleavage, including some protein component of the proteasome complex (PSM proteins). Cluster c3 express coding genes involved in cell migration, and underexpress PCGs involved in glucose metabolism. Cluster c4 contain overexpressed PCGs involved in epidermal development, including late cornified envelope (LCE) cluster genes and small proline-rich protein (SPRR) genes. Both LCE and SPRR proteins are expressed in terminally differentiated stratified epithelia, such as skin or head and neck mucosa. Underexpressed coding genes in c4 are involved in protein translation, similarly to underexpressed in c5. Cluster c5 has many overexpressed PCGs involved in DNA replication and cell cycle processes, RNA splicing or transcription. Overall, the results highlights important differences between lncRNA clusters in terms on predicted functions, somehow validating the unsupervised clustering and Pearson correlation analysis performed. Whether the surrogate lncRNAs of each cluster are also involved in these biological functions remains to be demonstrated.

**Figure 4.**
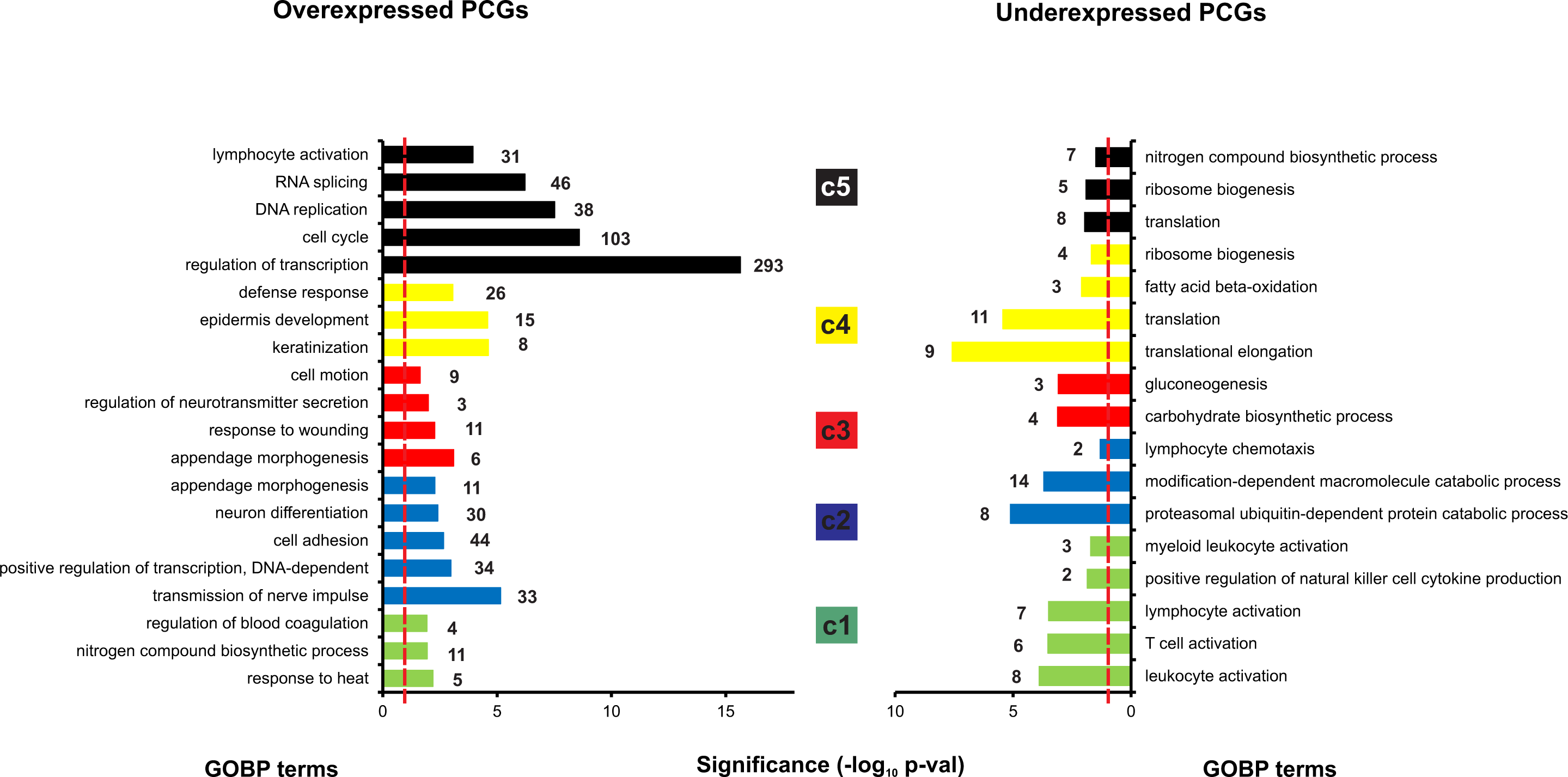
GOBP terms selected by cluster. Gene Ontology analysis was done using DAVID v6.7 web tool (see Methods). Overexpressed or a underexpressed PCGs are analyzed per cluster, and representative GO Biological Processes (GOBP) terms shown. Numbers at the end of each bar represent genes belonging to each GOBP term. Dashed red line denotes threshold for significant enrichment (p-val<0.05).

### Clinical and pathological features in lncRNA groups

We wanted to investigate whether our lncRNA clusters could display different clinical and pathological characteristics, possibly providing a better understanding of the patient’s tumor behavior or response to current therapies. We analyze whether patient sample clusters display follow-up differences in recurrence or in death, after treatment. Therefore, we performed Kaplan-Meier survival plots for both end-points (recurrence or overall survival) and the 5 clusters. Results show significant differences in overall survival at 5 years (p-val=0.0065, log-rank test) as previously described [16], with c5 patients having the highest probabilities of survival (Fig. 5A). In addition, patients within c1 and c2 clusters exhibit the lowest survival. No significant statistical differences were found, however, in recurrence after treatment (Supplementary Fig. 1).

**Figure 5.**
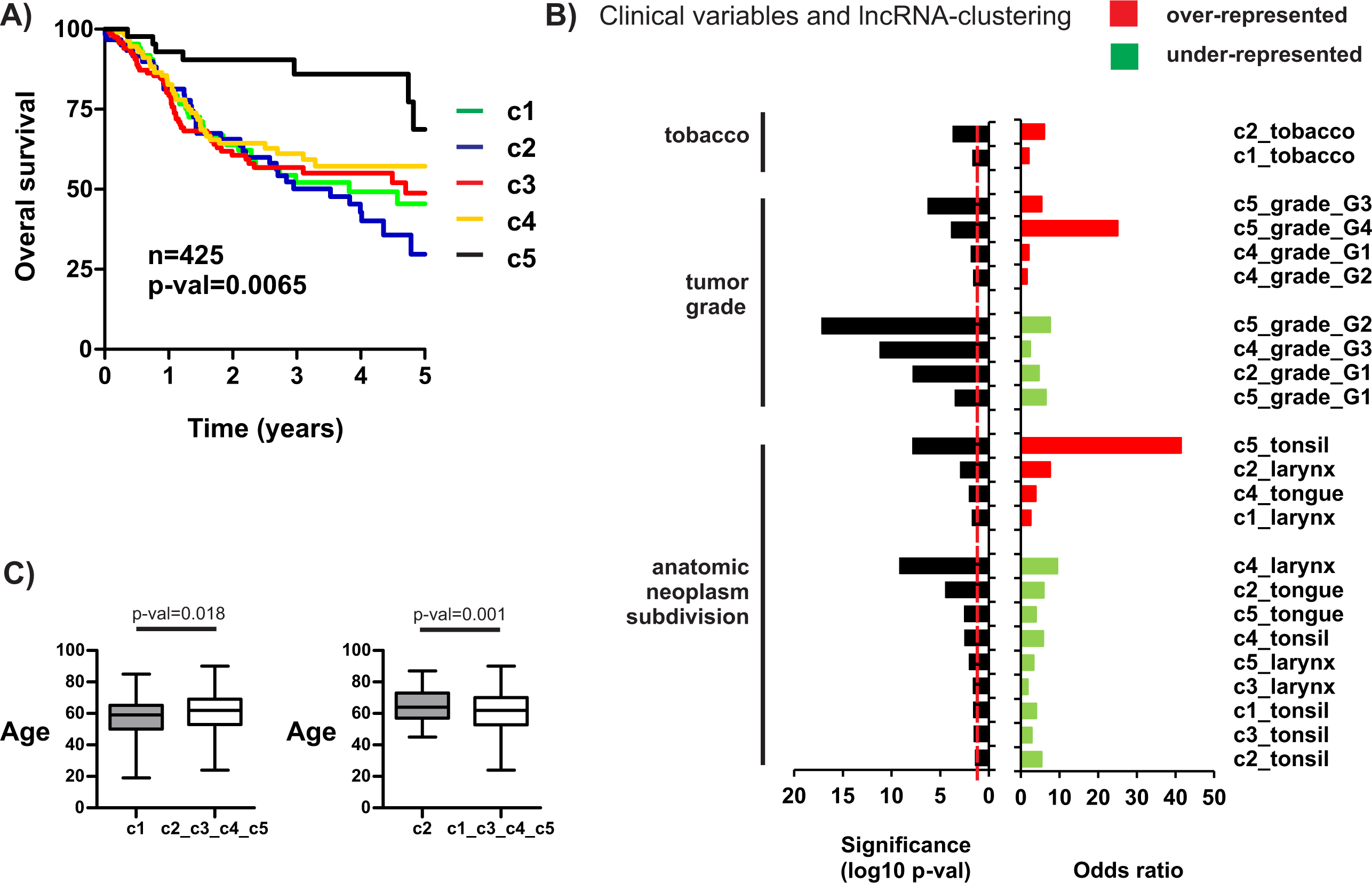
LncRNA-based clustering and clinical parameters. **A)** Kaplan-Meier plot of TCGA-HNSCC patients stratified by lncRNA clusters using overall survival as end-point. P-val was calculated with the log-rank test. n: number of patients with available follow-up information. **B)** Significant associations between lncRNA clusters and various clinical parameters, including tobacco, histologic grade, and sub-anatomical location. Odds ratios and significance values were calculated using Fisher’s exact test. **C)** Box plots of patient age for significant differences between c1 or c2 patients. P-values were calculated using Ttest.

Also, we examined tobacco smoking history, and found more smokers in c1 and c2 than expected by chance (p-val=0.02 and 0.0002, respectively, upon Fisher’s test) (Fig 5B). No differences were found in alcohol consumption between patient clusters. Interestingly, we show that tumor histology grade (G1, G2, G3, or G4) is associated with lncRNA clustering (p-val=6.58×10^−10^, Chi-square test). More specifically, we found that c5 tumors have frequently poorly differentiated and undifferentiated histology (G3 and G4, respectively) compared with the other clusters, while c4 have more differentiated or moderately differentiated tumors (G1 or G2, respectively). Furthermore, sub-anatomic location of the tumors and cluster subdivision were significantly associated (p-val=5.1×10^−41^, Chi-square test), so that c1 and c2 tumors are frequently found in larynx, c4 in tongue, and c5 in tonsil (Fig. 5B). Interestingly, 4 lncRNA genes from c1 and 10 from c2 have been shown to be overexpressed in larynx (Supplementary Table 6) [20]. Finally, we found that c1 patients are younger that the remaining patients (58.2 versus 61.6 years, p-val=0.018 after T-test), and c2 patients older (65 versus 60.2 years, p-val=0.001 after T-test) (Fig. 5C).

In summary, lncRNA clusterization is associated with important molecular aberrations and clinicopathologic features, providing important genomic and phenotypic relationships. Table 1 summarizes the main associated features of each lncRNA cluster.

**Table 1.** Summary of significant associations with lncRNA simple clusters

| **lncRNA cluster** | **c1** | **c2** | **c3** | **c4** | **c5** |
| --- | --- | --- | --- | --- | --- |
| HPV infection | negative | negative | negative | negative | positive |
| mutation ENRICHED | NSD1<br>TP53 | TP53<br>KMT2D |  | CASP8<br>CDKN2A<br>NOTCH1 |  |
| mutation DEPLETED | CASP8 | NOTCH1 | NSD1 | KMT2D<br>NSD1 | TP53<br>CDKN2A<br>FAT1 |
| lncRNAs UP | 91 | 275 | 9 | 13 | 245 |
| PCGs UP | 293 | 792 | 133 | 389 | 1485 |
| PCGs DOWN | 80 | 132 | 33 | 104 | 150 |
| age | younger | older |  |  |  |
| tobacco | smokers | smokers |  |  |  |
| tumor grade ENRICHED |  |  |  | G1, G2 | G3, G4 |
| tumor grade DEPLETED |  | G1 |  | G3 | G1, G2 |
| anatomic location ENRICHED | larynx | larynx |  | tongue | tonsil |
| anatomic location DEPLETED | tonsil | tongue<br>tonsil | larynx<br>tonsil | larynx<br>tonsil | tongue<br>larynx |

## Discussion

The recent implication of lncRNAs in many biological functions has established a new scenario to better understand complex processes like cancer [12, 13]. Traditional genomic characterizations have produced minimal improvements in patient clinical outcome, with a mortality rate of 50%. Therefore lncRNAs may represent a promising field to discover novel diagnostic and therapeutic strategies. Here we analyzed the lncRNA expression patterns of HNSCC from 426 TCGA primary tumor samples to generate insights into the landscape of lncRNAs in HNSCC.

Interestingly, we discover 5 clusters of samples based in lncRNA expression with significant differences in molecular aberrations and clinicopathologic features. LncRNA-clustering resembles clustering based on DNA-methylation and mRNA features. DNA-methylation is a chemical modification of genomic DNA by the addition of a methyl group (-CH3) to the cytosine or adenine DNA nucleotides. Typical DNA methylation occurs in a CpG dinucleotide context, where predominantly CpG sites are methylated in the genome. Most of the CpG clusters, known as CpG islands, occur near transcriptional start sites (TSSs) where they are predominantly un-methylated. The establishment and maintenance of methylation patterns resulting in modulation of gene expression is one of the key steps in epigenetic regulation during normal developmental programs. The significant overlapping between lncRNA and DNA-methylation clustering might be due to frequent methylation of CpG clusters closed to TSSs of lncRNAs altering their landscape. However, we cannot discard cross-regulation between DNA-methylation and specific lncRNAs, such as has been described for H19 [32] or ecCEBPA [33] recently. Interestingly, c1 contains almost all “hypo-methylated” samples from the DNA methylation clusters (Fig. 1A), concomitant with enrichment in NSD1 inactivating mutations. Similar mutations have also been found in squamous cell carcinoma of the skin [34]. This correlation between NSD1 mutations and the “hypo-methylated” samples has been previously reported in the TCGA HNSCC cohort [5, 35]. NSD1 is a histone 3 Lys 36 (H3K36) methyltransferase similar to SETD2, which is frequently mutated in the clear cell variant of renal cell carcinoma, and associated with DNA hypomethylation [36]. Moreover, a DNA hypomethylation signature has been reported in Sotos syndrome, a monogenic disorder defined by germline NSD1 mutations [37]. Several reports showed that H3K36 methylation is linked to the binding of de novo DNA methyltransferases (DNMT3A and DNMT3B) [38, 39]. Therefore, the recruitment of DNMT3A and DNMT3B could be impaired in NSD1 mutant tumors, leading to the global DNA hypomethylation observed in the c1 cluster. Whether this DNA hypomethylation is affecting c1 lncRNA expression regulation warrants further investigation. TP53 mutations have been associated with poor survival in many cancer types [40]. We believe that the aggressiveness of the disease in c1 and c2 patients might be, at list partially, explained by the increased TP53 mutation frequency. The reverse would apply for c5, almost depleted on mutations, and having the best overall survival of all clusters.

In relation with other clinical parameters analyzed, the anatomical location was the most significantly associated with lncRNA clustering, with c1 and c2 in larynx, c4 in tongue, or c5 in tonsil. This finding might be explained by the already known high tissue specificity of lncRNA expression [41], and highlights an important issue: HNSCC as a group includes different tumor locations and different etiologies. Therefore, we consider that some of the surrogate lncRNAs in the clusters might be specific for sub-anatomical locations in the head and neck region, and may help to analyze location-specific associations between molecular and phenotype features. For example, c5 overexpressed coding-genes are enriched in proteins involved in “lymphocyte activation”, a process that occurs in the tonsils, which is an immune defense organ in the aerodigestive tract constituting the first line of defense against ingested or inhaled foreign pathogens. Naturally, both B- and T-cell activation occurs in the tonsils after the uptake of antigens produced by pathogens by specialized antigen capture cells called M cells. Whether the enrichment in “lymphocyte activation” found in c5 is due to tonsil-dependent functions or to a specific immune response to HPV-positive tumors remains to be ascertained.

Between the PCGs underexpressed in c1, we found programmed death-ligand 1 (PD-L1; also called CD274). PD-L1, which is expressed on many cancer and immune cells, plays an important part in blocking the ‘cancer immunity cycle’ by binding programmed death-1 (PD-1) and B7.1 (CD80), both of which are negative regulators of T-lymphocyte activation [42]. Binding of PD-L1 to its receptors suppresses T-cell migration, proliferation and secretion of cytotoxic mediators, and restricts tumor cell killing. The PD-L1-PD-1 axis protects the host from overactive T-effector cells not only in cancer but also during microbial infections. Blocking PD-L1 should therefore enhance anticancer immunity and successful treatment of many patients with advanced cancer using antibodies against PD-L1 has been demonstrated, including HNSCC [43]. Thus, the Food and Drug Administration (FDA) recently approved pembrolizumab, an antibody inhibiting PD-L1, for the treatment of some patients with recurrent or metastatic HNSCC that has continued to progress despite standard-of-care treatment with chemotherapy. However, little is known about predictive factors of efficacy of such therapies, although some recent reports described that across multiple cancer types, responses are observed in patients with tumors expressing high levels of PD-L1, especially when PD-L1 was expressed by tumor-infiltrating immune cells. Therefore, it is tempting to speculate that c1 primary tumors would be less sensitive to anti-PD-L1 therapies than the tumors from the other clusters. In addition, some of the c1 lncRNAs might be involved in this PD-L1-PD-1 ‘cancer immunity cycle’, and could be subject of future investigations.

Coding genes involved in synaptic transmission and neuron differentiation are coexpressed with surrogate lncRNAs within cluster c2. Perineural invasion occurs in an important proportion of HNSCC samples. We found no significant higher frequency of perineural invasion in c2 samples compared with the remaining samples (Fisher’s exact test, p-val=0.5). Therefore, it is tempting to speculate that c2 samples might display some cellular plasticity from epithelial towards neuroendocrine lineage, as HNSCC tumors with neuroendocrine phenotype has been described [44]. Recently, neuroendocrine lineage plasticity enabled by the loss of TP53 and RB1 function was shown in prostate cancer, mediated by increased expression of the reprogramming transcription factor SOX2 [45]. Interestingly, c2 display higher frequency of TP53 mutations (Fig. 2C and Table 1). Possibly, lncRNAs within c2 might be involved in mechanisms regulating epithelial to neuroendocrine reprograming.

Many overexpressed PCGs in c5 are related to “cell cycle” processes, suggesting that c5 carcinomas are more proliferative. This finding is in line with the oncogenic activities of HPV E7 oncoprotein, which binds and induces protein degradation of cell cycle regulators such as the retinoblastoma protein family, including pRb (RB1), p107 (RBL1) or p130 (RBL2) [46]. Degradation of these proteins, mainly pRb, allow E2F transcription factors to induce expression of genes involved in cell cycle progression [47], such as cyclins and cyclin-dependent kinases, or proteins involved in DNA replication, and mitotic division. In addition, E2F1 gene amplification was found in HPV positive HNSCC tumors from the TCGA cohort [5], which correlates with a molecular profile of cell cycle deregulation. Therefore, some c5 lncRNAs might also be involved in these processes, and possibly in the E7-pRb-E2F axis of cell cycle deregulation of HPV-infected tumors. Tumors from c4 overexpresses PCGs involved in epidermal differentiation, such as LCE cluster SPRR genes, normally involved in terminal epithelial differentiation. This result might be in line with the high frequency of differentiated carcinomas in c4 (histologic grades G1 and G2) (Fig. 5 and Table 1), whereby G3 tumors are less frequent. Therefore, some of the c4 surrogate lncRNAs could be involved in epithelial differentiation.

## Conclusions

We present the first comprehensive clustering analysis of HNSCC based on lncRNA expression performed to date. The results allow selection of surrogate lncRNA genes of 5 distinct tumor groups, propose possible functions associated with them, as well as phenotypic and clinico-pathology features that may be consequence, in part, of their activities.

## Declarations

### Ethics approval and consent to participate

Not applicable

### Consent for publication

Not applicable.

### Availability of data and materials

The datasets generated and/or analyzed during the current study are available in the TANRIC (http://ibl.mdanderson.org/tanric/_design/basic/index.html), TCGA (https://tcga-data.nci.nih.gov/docs/publications/hnsc_2014/) repositories, and webtools as cBioPortal (http://www.cbioportal.org/) and UCSC Xena (http://xena.ucsc.edu/).

All data generated or analyzed during this study are included in this published article [and its supplementary information files].

### Authors’ contributions

PGdeL, APG and RGE perfomed the genomic and statistical analyses. PGdeL and RGE wrote the manuscript. PGdeL, JMP and RGE interpreted the results. All authors read and approved the final manuscript.

## Acknowledgements

Not applicable

---

**Supplementary Figure 1.** Kaplan-Meier plot of TCGA-HNSCC patients stratified by lncRNA clusters using recurrence and end-point. P-val was calculated with the log-rank test. n: number of patients with available follow-up information.

